# Outer membrane vesicles derived from *Klebsiella pneumoniae* are a driving force for horizontal gene transfer

**DOI:** 10.1101/2021.07.06.451238

**Authors:** Federica Dell’Annunziata, Carmela Dell’Aversana, Nunzianna Doti, Giuliana Donadio, Fabrizio Dal Piaz, Viviana Izzo, Anna De Filippis, Marilena Galdiero, Lucia Altucci, Giovanni Boccia, Massimiliano Galdiero, Veronica Folliero, Gianluigi Franci

## Abstract

Gram-negative bacteria release outer membrane vesicles (OMVs) into the extracellular environment. Recent studies recognized these vesicles as vectors to horizontal gene transfer, however the parameters that mediate OMVs transfer within bacterial communities remain unclear. The present study highlights for the first time the transfer of plasmids containing resistance genes via OMVs derived from *Klebsiella pneumoniae* (*K. pneumoniae*). This mechanism confers DNA protection and it is plasmid copy number dependent with a ratio of 3.6 time among high copy-number plasmid (pGR) versus low copy number plasmid (PRM) and the transformation efficiency was 3.6 times greater. Therefore, the DNA amount in the vesicular lumen and the efficacy of horizontal gene transfer was strictly dependent on the identity of the plasmid. Moreover, the role of *K. pneumoniae*-OMVs in interspecies transfer was described. The transfer ability was not related to the phylogenetic characteristics between the donor and the recipient species. *K. pneumoniae*-OMVs transferred plasmid to *Escherichia coli*, *Salmonella enterica*, *Pseudomonas aeruginosa* and *Burkholderia cepacia*. These findings address the pivotal role of *K. pneumoniae*-OMVs as vectors for antimicrobial resistance genes spread, contributing to the development of antibiotic resistance in the microbial communities.

**Author summary:** *K. pneumoniae* is an important opportunistic pathogen that affects several host districts, in particular respiratory and urinary tracts. Hospital-acquired *K. pneumoniae* infections lead to a 50% mortality rate correlated with rapid acquisition of antibiotic resistance. Currently, the increasing rate of antibiotic resistance among *K. pneumoniae* isolates is a major concern worldwide. The spread of multidrug-resistant *K. pneumoniae* strains renders current therapeutic options ineffective. Like all Gram-negative bacteria, *K. pneumoniae* secretes OMVs. OMVs are spherical structures, with a diameter between 50-250 nm, originating from the outer membrane. OMVs biogenesis allows bacteria to interact with the external environment, increasing bacterial survival under stressful conditions and regulating microbial interactions within bacterial communities. Few evidence recognized OMVs as vectors for horizontal gene transfer, contributing to the spread of resistance. In this scenario, the present study examines the potential role of *K. pneumoniae*-OMVs in inter- and intra-species diffusion of ß-lactam resistance.

## Introduction

Horizontal gene transfer (HGT) represents the main source of genetic material transfer among microorganisms [1]. Indeed, HGT provides a driving force for bacterial evolution, increasing bacterial survival, adjustment rate in the harshest environments and pathogenicity [2–4]. Current knowledge of HGT is based on three widely described mechanisms for the exchange of genetic material between bacteria: transformation, conjugation and transduction [5–7].Transformation involves the natural uptake of naked DNA from an extracellular environment; this phenomenon occurs when cells are in a physiological state of competence, regulated by 20-50 proteins [8,9]. Conjugation is a DNA transfer mechanism through the sexual pilus and requires cell-to-cell contact [10]. Conjugative systems are frequently associated with plasmid transfer [11]. Transduction entails the transfer of DNA between bacteria through the bacteriophage infections [12]. The recombinant phage particle can contain up to 100 kilobases of DNA and the infection is limited to host specificity [13]. Recently, several studied reported that HGT processes is facilitated by Outer Membrane Vesicles (OMVs) [14–17].

OMVs are spherical nanostructures, 50–250 nm in diameter, released naturally and constitutively by Gram-negative bacteria during their growth [18]. OMVs originate from the outer membrane (OM) and include in the vesicular lumen lipopolysaccharide, peptidoglycan, phospholipids, genetic material (DNA and RNA) and periplasmic and cytoplasmic protein components, during their biogenesis [19]. Although many aspects of vesicular biogenesis and regulation of their composition remain unclear, the biological functions associated with OMVs release were extensively described [20,21]. These vesicles play a key role in the bacteria-environment, bacteria-bacteria and bacteria-host interactions [22,23]. OMVs are recognized for their role in nutrient acquisition, response to stress, biofilm formation and toxins release, adhesion and virulence factors and in host defense system evasion [24]. OMVs role in HGT was reported in *E. coli*, *Acinetobacter baumannii*, *Acinetobacter baylyi*, *Porphyromonas gingivalis*, *P. aeruginosa* and *Thermus thermophilus* [25–28]. Yaron et al. demonstrated the transfer of virulence genes through *E. coli*-OMVs between bacteria of different species. Moreover, they proved that the genetic material was protected from digestion with DNase, confirming the packaging in the vesicular lumen [29]. OMVs derived from *A. baumannii* were also identified as vectors for antibiotic resistance gene transfer. In the study of Rumbo et al. plasmid-borne bla_OXA-24_ gene conferred carbapenems resistance to sensitive Acinetobacter strains [30]. These evidences highlight the potential OMVs contribution to the spread of virulence and antibiotic resistance which represents, to date, a serious risk to human health.

In this scenario, *K. pneumoniae* represents one of the most worrying pathogens involved in nosocomial infections [31]. The constant antibiotics treatment induces selective pressures, causing the evolution of multidrug-resistant (MDR) bacteria [32]. Our previous studies demonstrated that OMVs derived from *K. pneumoniae* play a crucial role in the microorganism-host interaction, modulating miRNAs genetic transcription and influencing the inflammatory response [14,33]. Currently, no study showed the role of *K. pneumoniae*-OMVs as a carrier for HGT, allowing the transport of genetic material and the spread of resistance genes. Therefore, this study demonstrates, for the first time, *K. pneumoniae*-OMVs HGT role. We investigated OMVs contribution in the genetic material cargo and in intra and inter-species transfer. After, we collected evidence to demonstrate that plasmid copy number (PCN) might play an important role in the biogenesis, cargo and in the HGT mechanisms. Finally, we verified OMVs stability over time and whether storage conditions might influence gene transfer.

## Results

### Characterization of isolated *K. pneumoniae*-OMVs

To purify the OMVs derived from *K. pneumoniae*-pGR and *K. pneumoniae*-PRM, bacteria were grown in LB supplemented with ampicillin up to the late logarithmic-phase of the bacterial growth curve (see materials and methods for specifications). Vesicles were collected from culture supernatants and characterized in terms of morphology, size and polydispersity index (PDI). Purified OMVs appeared at TEM as electron-dense particles, with uniform spherical morphology (figure 1, panel A and B). No bacterial contaminant was visualized, demonstrating the total sterility of the vesicular suspensions used. Dynamic light scattering (DLS) analysis showed that OMVs derived from *K. pneumoniae*-pGR measured a size of 113.8 ± 53.7 nm and were characterized by a slightly heterogeneous size distribution, represented by the PDI of 0.223 (Figure 1 C). OMVs purified from *K. pneumoniae*-PRM showed a reduction in size and higher vesicular populations homogeneity, recording a size of 94.13 ± 41.10 nm and a PDI of 0.191 (Figure 1 D). Purified OMVs were also characterized based on protein profile. The total vesicular proteins were extracted from *K. pneumoniae*-OMVs via lysis buffer and then quantified by Bradford assay. The protein amount was 35.77 mg and 30.00 mg for *K. pneumoniae*-pGR and *K. pneumoniae*-PRM, respectively, obtained from 600 mL of culture supernatant. Five micrograms of protein were loaded on 10% SDS-PAGE and the gel was stained with Blue Coomassie (Supplementary figure 1). The corresponding bands were excised and subjected to in-situ digestion protocols. Peptides were analyzed by high resolution nanoLC-MS / MS. Mass spectra analysis allowed identifying with high confidence proteins common to both purified OMVs samples (Supplementary table 1). These proteins were classified according to the subcellular localization site and biological function (Figures 2, A and B). The vesicles contained 14 membrane-associated proteins (28.57%), 3 periplasmic proteins (6.12 %) and 32 cytosolic proteins (65.31%). In addition, 3 DNA-binding proteins (6.12%) were identified among the proteins annotated for their binding function. Twenty-one enzymes were revealed, including 3 oxidoreductases (14.29%), 2 transferases (9.52%), 1 aminopeptidase (4.76%), 5 lyases (23.81%), 2 isomerases (9.52%) and 8 ligases (38.10%), (Figure 2 C). To confirm accurate reliability and reproducibility of data, three independent OMVs purifications were performed and analyzed for each strain.

**Figure 1.**
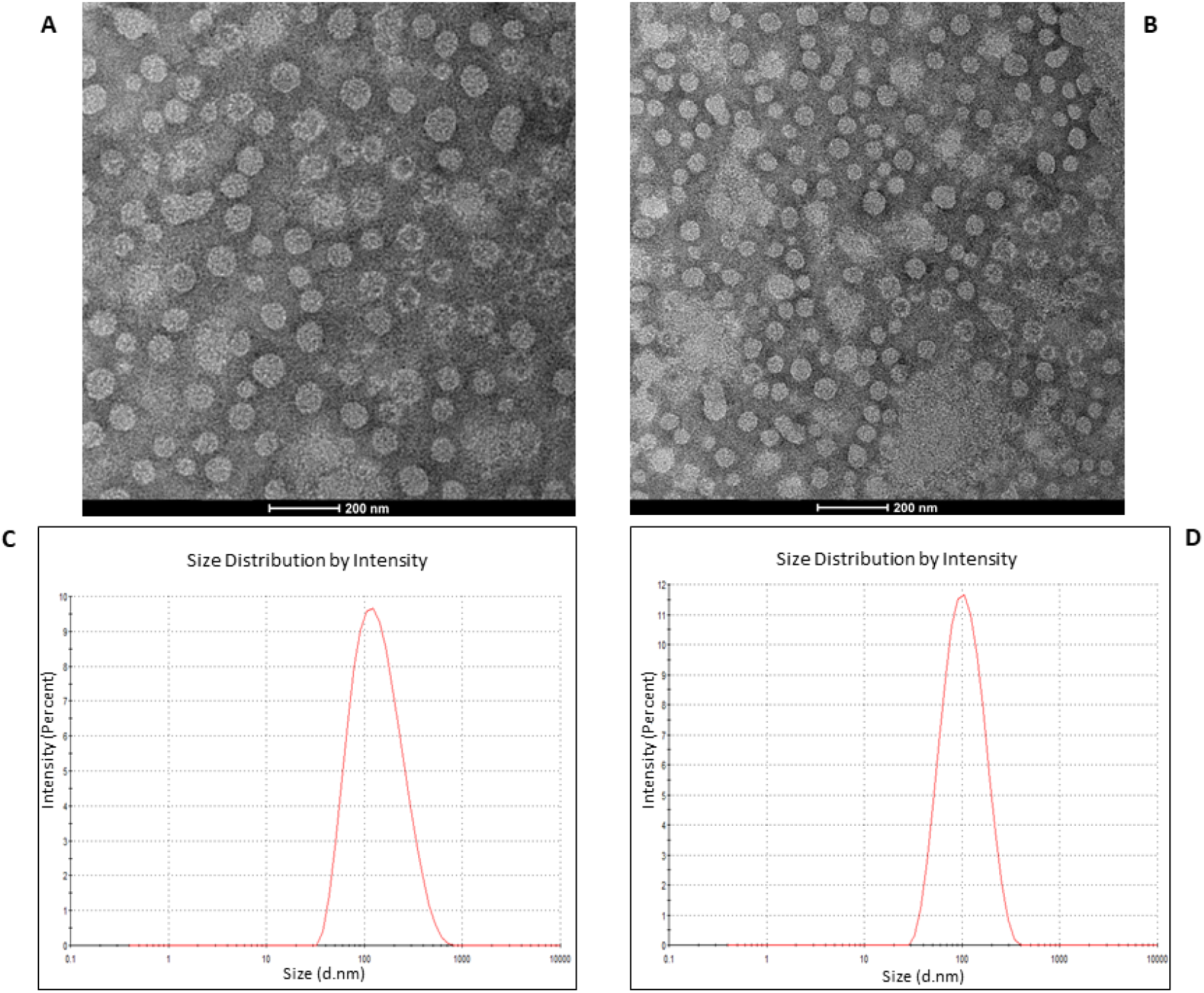
TEM of OMVs purified from *K. pneumoniae*-pGR (A) and *K. pneumoniae*-PRM (B) (scale bar = 200 nm). DLS intensity-weighed distribution of OMVs derived from *K. pneumoniae*-pGR (C) and *K. pneumoniae*-PRM (D).

**Figure 2.**
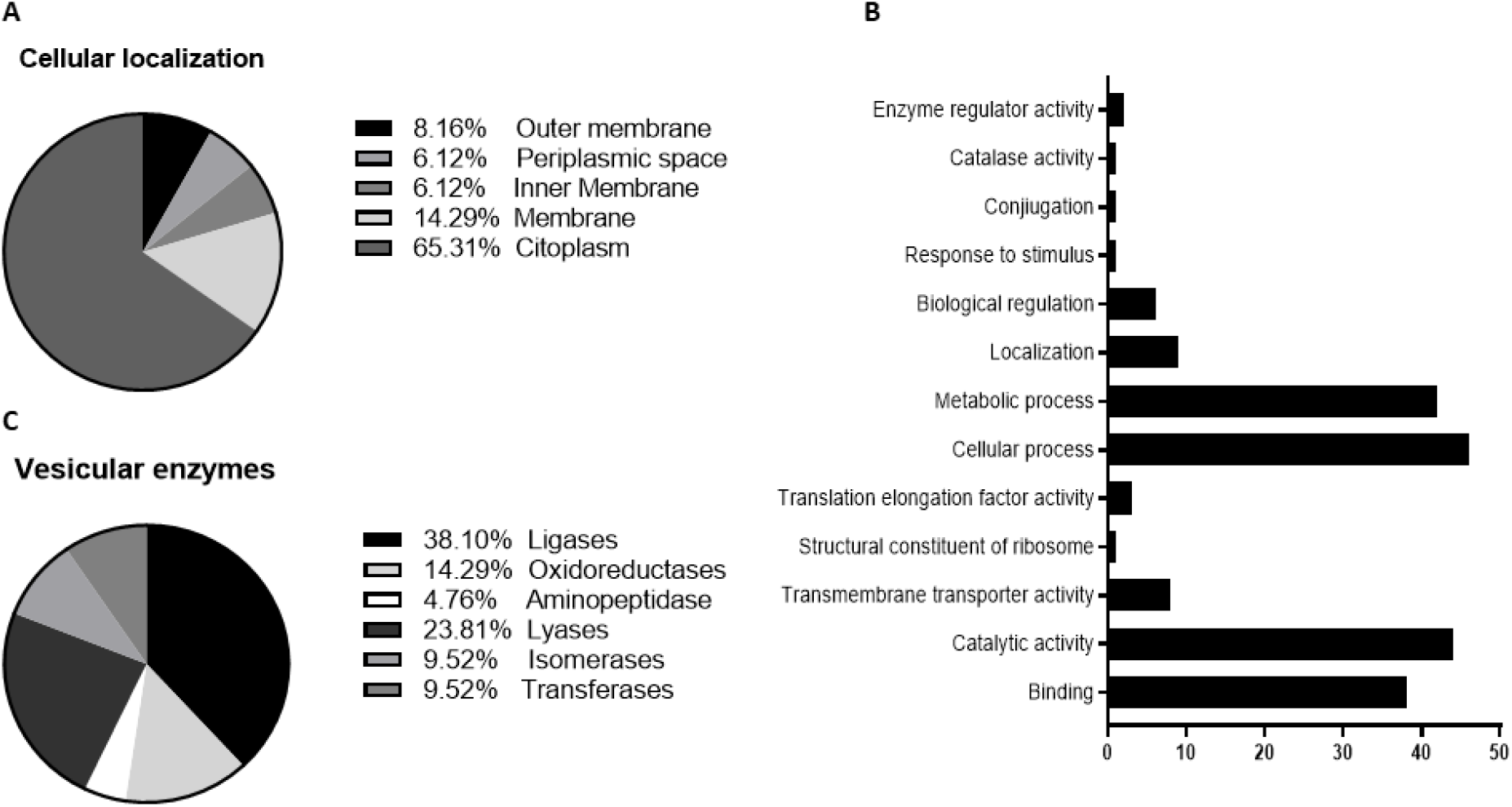
Classification of cellular localization (A), functional annotation (B) and enzymatic classes (C) of protein extracted from *K. pneumoniae*-OMVs.

### DNA packaging in *K. pneumoniae* OMVs

The propensity of *K. pneumoniae* OMVs to incorporate genetic material during the biogenesis process was evaluated by transforming bacteria with pGR and PRM plasmids. *K. pneumoniae*-pGR and *K. pneumoniae*-PRM were grown on LB supplemented with 100 μg mL^-1^ of ampicillin for selection of transformants. Plasmid DNA extraction and enzymatic digestion profile confirmed the plasmids presence in the bacterial strains (Supplemental figure 2 A and B). The presence of pGR and PRM plasmids in *K. pneumoniae*-OMVs was evaluated by absolute qPCR. To demonstrate that DNA was present in the vesicular lumen and protected from the extracellular nucleases action, qPCR was performed using OMVs samples either treated or untreated with DNase. In untreated OMVs, plasmid concentration was 18.91 ± 0.53 and 14.78 ± 0.91 ng DNA/μg OMVs, for pGR and PRM, respectively. In OMVs treated with DNase before vesicular lysis, pGR recorded a higher loading density, measuring 10.4 ± 0.05 ng DNA /μg OMVs, which corresponded approximately to 1.9 × 10^9^ PCN /μg OMVs. Otherwise, PRM measured a plasmid concentration of 3.08 ± 0.62 ng DNA /μg OMVs, corresponding to 9.6 × 10^8^ PCN /μg OMVs (Figure 3).

**Figure 3.**
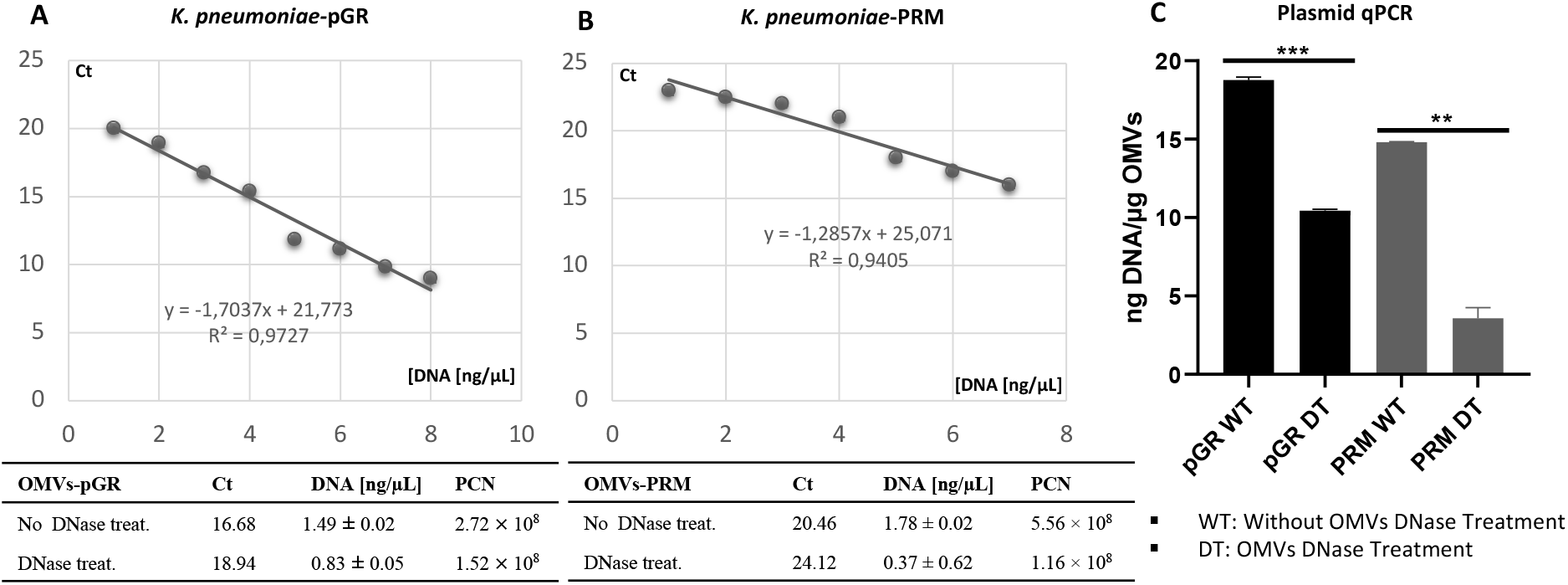
Determination of PCN in OMVs, using quantitative PCR standard curves. The standard curves were generated by qPCR of the purified pGR (A) and PRM (B) plasmids. Histogram Graph of PGR and PRM cargo efficiency, before and after DNase treatment (C) (*P value* < *0.05*).

### OMVs mediate the plasmid intra-specie transfer

Transformation experiments were performed by isolating OMVs from *K. pneumoniae*-pGR and *K. pneumoniae*-PRM. *K. pneumoniae* ATCC recipient cells were incubated with 10 μg of OMVs derived from *K. pneumoniae*-pGR and *K. pneumoniae*-PRM. After 24 hours, treated cells were plated on LB-ampicillin agar to detect the plasmid resistance marker in the recipient bacteria. OMVs purified from *K. pneumoniae*-pGR induced a transformation efficiency of 2.8 ± 0.1 x 10^4^ CFU / μg. HGT mediated by *K. pneumoniae*-PRM-OMVs occurred with a transformation efficiency of 7.8 ± 0.9 x 10^3^ CFU / μg. In both conditions, no plasmid acquisition occurred when recipient cells were incubated with free plasmid (Figure 4 A-H). Therefore, HGT via OMVs derived from *K. pneumoniae*-pGR was 3.6 times more efficient than the *K. pneumoniae*-PRM OMVs transfer, with the same vesicular concentration (Figure 4 I). Colony-PCR was used to confirm that resistant acquisition. PGR and PRM were determined by amplifying a region of the β-lactamase gene and the amplicon was visualized by agarose gel electrophoresis (Figure 5 A and B). PCR analysis showed that all grown and selected colonies on LB-ampicillin plates contained pGR and PRM plasmids. Pre-transformation colonies of *K. pneumoniae* did not show amplification, demonstrating the absence of resistance. Amplification of the 16S ribosomal gene region was used as a housekeeping control.

**Figure 4.**
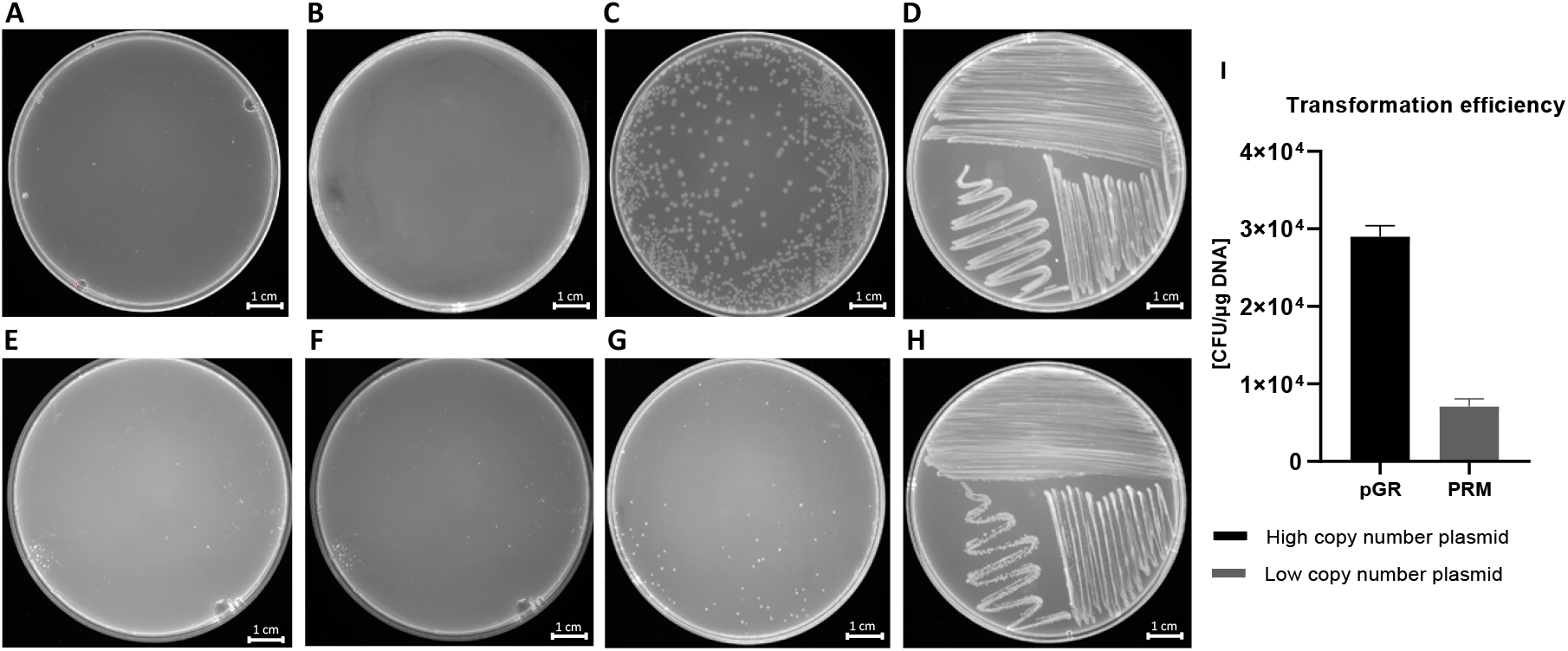
HGT via OMVs derived from *K. pneumoniae*-pGR. Untreated cells (A) and cells treated with free plasmid (B) did not record transformants. *K. pneumoniae* ATCC treated with 10 μg of OMVs (C) and bacteria control on LB-plates (D). HGT-OMVs from *K. pneumoniae*-PRM. Transformation in untreated (E) and treated with free plasmid (F) bacteria did not occurred. *K. pneumoniae* ATCC incubated with 10 μg of OMVs (G) and bacteria control on LB-plates (H). Intra-species HGT efficiency via OMVs purified from *K. pneumoniae*-pGR and *K. pneumoniae*-PRM.

**Figure 5.**
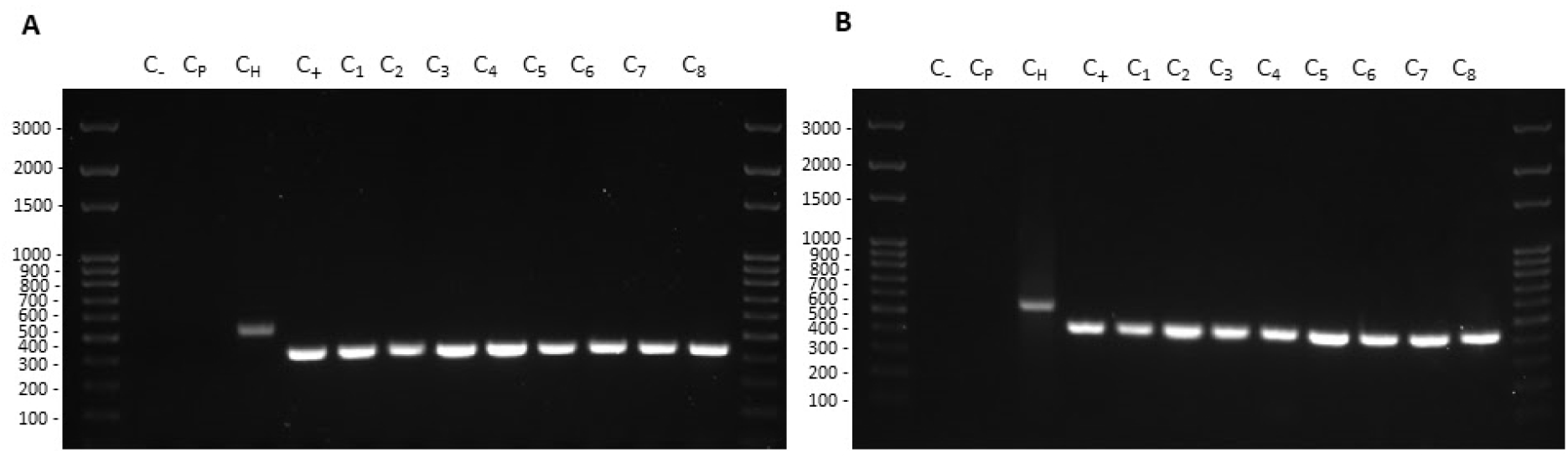
Colony-PCR from recipient cells treated with *K. pneumoniae* pGR (A) and *K. pneumoniae* PRM (B) OMVs. DNA gel showed PCR products with expected lengths: ß-lactamase product~ 424 bp (C_1-8_), ribosomal 16S product ~ 550bp (C_H_). Control water (C_-_) and untreated bacteria (C_p_) did not show amplification.

### OMVs induce the generalized resistance spread

The OMVs potential to transfer genetic material between different microbial species was evaluated. Five recipient bacterial species were selected based on taxonomic differences (Figure 6A). Cultures of *K. pneumoniae*, *E. coli*, *S. enterica*, *P. aeruginosa* and *B. cepacia* were treated with *K. pneumoniae*-pGR OMVs. After 24 hours of incubation, recipient cells were plated on LB-ampicillin agar plates and counted to define the transformation efficiency. OMVs derived from *K. pneumoniae*-pGR transferred plasmid DNA with a transformation efficiency of 2.8 ± 0.1 x 10^4^, 1.7 ± 0.2 x 10^4^, 1.5 ± 0.9 x 10^4^, 1.6 ± 0.1 x 10^4^, 1.8 ± 0.8 x 10^4^ CFU / μg for *K. pneumoniae*, *E. coli*, *S. enterica*, *P. aeruginosa* and *B. cepacia*, respectively (Figure 6 B). Colonies of each recipient bacterial species were selected and subjected to PCR analysis to confirm the presence of the β-lactamase gene in the recipient species (Supplementary figure 3 A, B, C and D). Recipient cells incubated with free plasmid pGR and untreated cells did not acquire antibiotic resistance in any condition.

**Figure 6.**
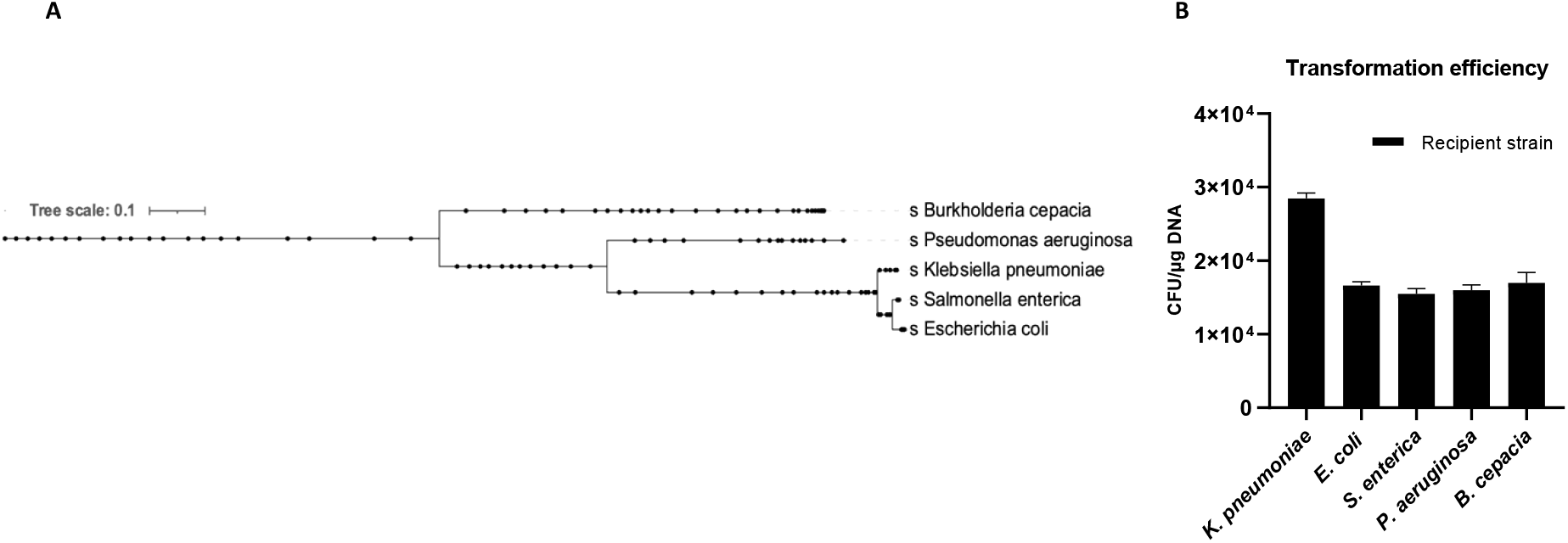
Phylogenetic relationship of the recipient species (A). *K. pneumoniae*-pGR OMVs inter-species transformation efficiency (B).

### OMVs stability over time

OMV-HGT experiments continued to evaluate transformation efficiency over time, by storing OMVs derived from *K. pneumoniae*-pGR at −20 ° C for 30 days and at +4° C for 7 days. A gradual reduction in transformation efficiency was observed using OMVs treated with DNase and stored at −20 ° C and +4° C for increasing periods of time. The HGT experiment was conducted using *K. pneumoniae* ATCC as a recipient cell. The maximum number of transforms was obtained with OMVs used after 10 days of storage, showing an efficiency of 2.5 ± 0.1 x 10^4^ CFU / μg. After 20 days, a reduction in efficiency was verified, recording 7.7 ± 0.9 x 10^3^ CFU / μg. On the 30th day of storage, a drastic decrease of transformants had occurred, registering 9.1 ± 0.12 x 10^2^ CFU / μg. The OMVs stability at 4 ° C showed a slight reduction over time. The recorded transformation efficiency was 1.7 ± 0.25 x 10^4^ and 8.0 ± 0.43 x 10^3^ CFU / μg, after 3 and 7 days of storage, respectively (Figure 7).

**Figure 7.**
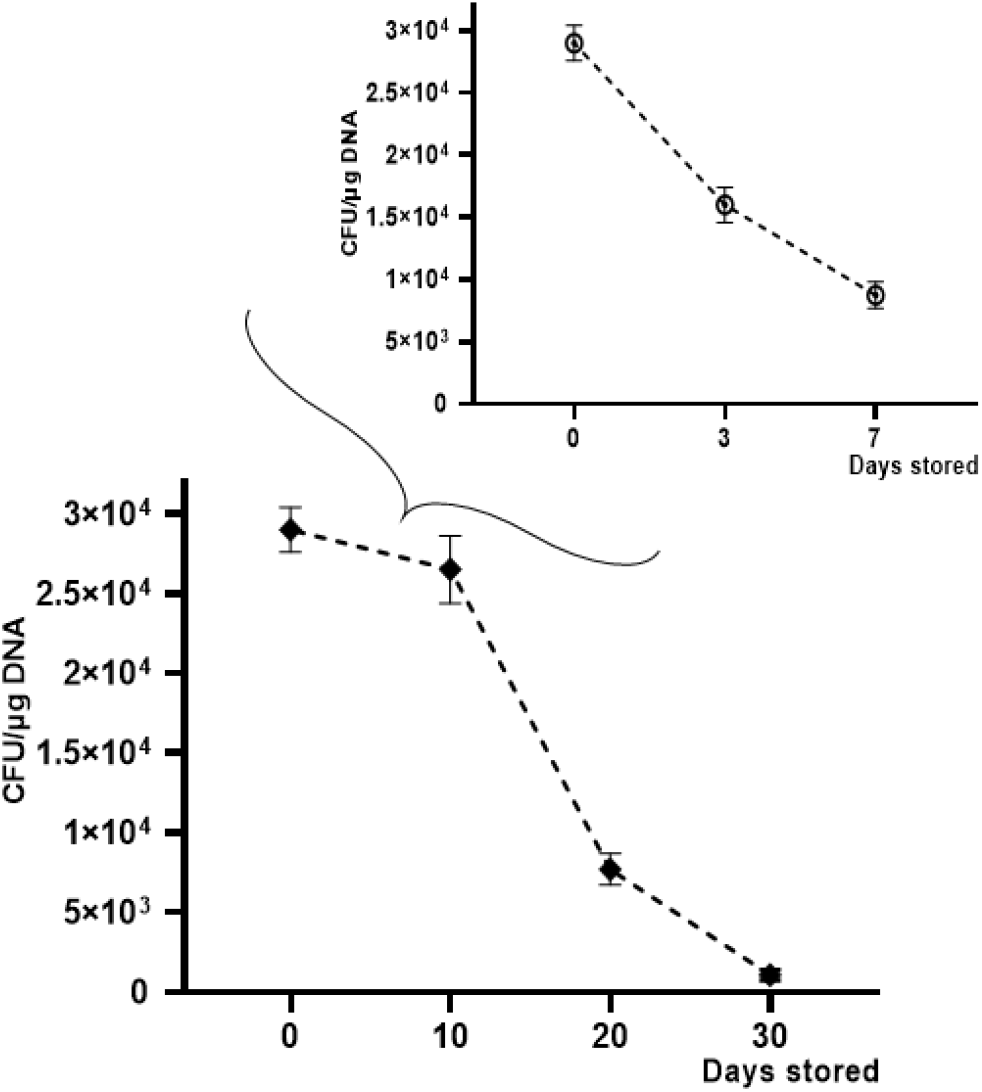
Transformation frequency of *K. pneumoniae*-pGR OMVs storage at −20 °C and + 4°C for 30 and 7 days, respectively.

### Diagnostic transformed strains characterization

The phenotypic effect correlate to the genotypic resistance detected by PCR analysis was evaluated through antibiotic susceptibility testing. Each bacterial strain, before and after treatment with OMVs derived from *K. pneumoniae* pGR, was examined. Concerning the susceptibility to β-lactams, the inhibition diameter measured before OMVs treatment were 14.5 ± 0.3, 14 ± 0.01, 18.3 ± 0.07, 25 ± 0.5, 20 ± 0.9 mm for *K. pneumoniae*, *E. coli*, *S. enterica*, *P. aeruginosa* and *B. cepacia* respectively. Inhibition zones recorded were associated with susceptible strains, in accordance with EUCAST guidelines. After the OMV-HGT, no inhibition area was identified for β-lactam antibiotics, demonstrating the acquisition of resistance. The inhibition area measured for ciprofloxacin was ≥ 30 mm, before and after OMVs treatment, in each bacterial species (Figure 8). The ciprofloxacin control was used to demonstrate that the acquired resistance was associated with the plasmid containing β-lactamase gene.

**Figure 8.**
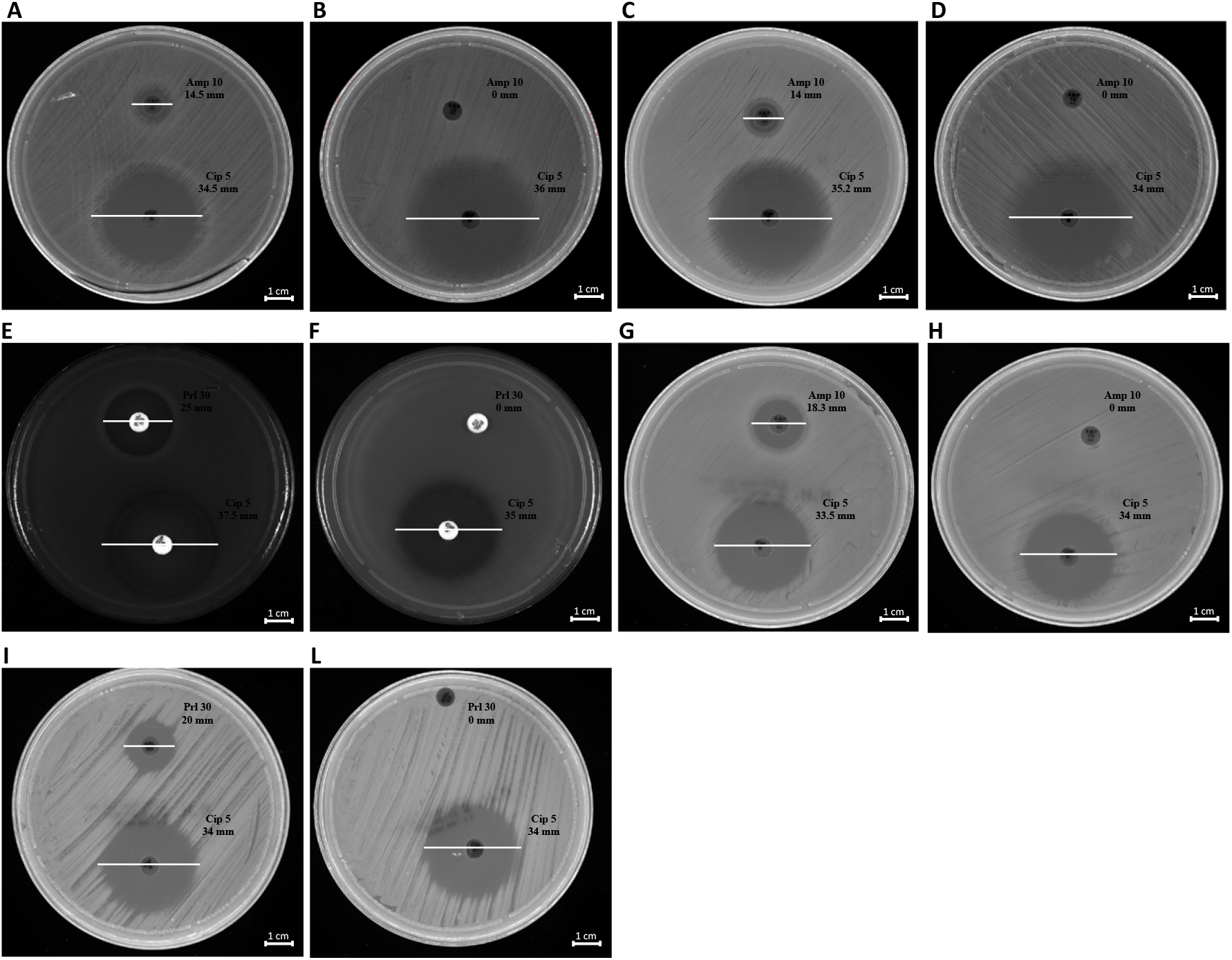
Antibiotic susceptibility of bacteria to ß-lactams. *K. pneumoniae* pre and post-OMVs treatment (A, B); *E. coli* pre and post-OMVs treatment (C, D); *P. aeruginosa* pre and post-OMVs treatment (E,F); *S. enterica* pre and post-OMVs treatment (G, H); *B. cepacia* pre and post-OMVs treatment (I, L).

## Discussion

Horizontal gene transfer plays an important role in promoting bacterial evolution, adaptation to environmental changes and acquisition of new metabolic capabilities [3]. Genetic pool modifications as a consequence of genetic transfer were observed in bacterial communities with high frequency rates, demonstrating the importance of this phenomenon for bacterial survival [34]. Currently, transformation, transduction and conjugation are considered the three canonical HGT mechanisms, contributing significantly to genetic diversity [35]. However, novel genetic material exchange events are under consideration and may be added to those currently known. Nowadays, the HGT mechanism should include the OMVs secretion by Gram-negative [36]. Previous studies reported that OMVs incorporated DNA into the lumen and transported it to recipient cells [37]. Currently, no studies assessed the ability of *K. pneumoniae* to exploit HGT via OMVs to spread antimicrobial resistance. However, multidrug-resistant *K. pneumoniae* is increasingly implicated in hospital-acquired infections, causing high morbidity and mortality. Improved understanding of *K. pneumoniae* mechanisms to resistance-genes spread is needed. Therefore, the focus of this research was the preliminary characterization of HGT mechanisms mediated by OMVs derived from *K. pneumoniae*.

Firstly, OMVs were isolated from *K. pneumoniae*-pGR and *K. pneumoniae*-PRM, respectively. TEM and DLS analysis revealed that the vesicles featured a spherical morphology, in accordance with our previously published data, but with a reduced diameter compared with the OMVs collected from *K. pneumoniae* ATCC [14]. The different vesicular size could be attributed to the antibiotic presence during bacterial growth. Indeed, Fulsundar et al. showed that antibiotic and environmental stresses determined a significant effect on the OMVs production, size and DNA content [17]. These evidences confirm that OMVs release is a physiologically controlled process, dependent on environmental factors. The proteomic characterization of OMVs derived from *K. pneumoniae*-pGR and *K. pneumoniae*-PRM identified more than 55 proteins, mainly from the outer membrane and the periplasmic space. Inner membrane and cytosolic proteins were also detected, demonstrating that, although the mechanism of inclusion is unclear, cytoplasmic components and portions of membrane were incorporated into OMVs during the biogenesis process. Finally, the presence of proteins capable of interacting with DNA could confirm the ability of OMVs to also incorporate genetic material. These results may suggest, in addition to the OMVs originating from outer membrane budding, the possible existence of another vesiculation pattern. Indeed, Cruez et al. showed in *S. vesiculosa* M7T, *N. gonorrhoeae*, *P. aeruginosa* PAO1 e *A. baumannii* AB41 the presence of vesicles containing a bilayer of membrane and highly electrodense cytoplasmic material. These vesicles were classified as outer-inner membrane vesicles (O-IMVs) [38]. The possible secretion of two vesicle types, OMVs and O-IMVs, could explain how DNA is incorporated in OMVs, since is not properly clarified. Currently, three models were proposed: i) the DNA present in the extracellular environment was internalized according with a mechanism similar to bacterial transformation; ii) DNA was transported through the inner membrane and the cell wall up to the periplasmic space, where it was included in the OMVs; iii) the DNA inclusion in the vesicles occurred through the secretion of O-IMV, which incorporate cytoplasmic components and DNA. The third model is the most accredited and supported by experimental evidence [20]. Although the DNA inclusion mechanism is not known with absolute certainty, our finding demonstrated that OMVs secreted by *K. pneumoniae* were involved in HGT, allowing the spread of resistance genes in microbial communities.

Contextually, our manuscript demonstrated that *K. pneumoniae* transferred genetic material, incorporating DNA within the OMVs and protecting it from the extracellular exonucleases action. The DNA in the vesicular lumen was transferred to the recipient cell by determining the acquisition of resistance genes present in the plasmid. The recipient cell *K. pneumoniae*, after contact with OMVs, acquired and expressed resistance to ampicillin, proving the OMVs ability to promote intraspecies HGT. Plasmid transfer did not occur when cells were incubated with free plasmid, suggesting that vesicles could represent a physiological mechanism that exceeds environmental limits (exonuclease degradation, dilution of gene material, long-distance transfer, etc.) and associated with the donor / recipient cell (state of competence, high vesicle-OM affinity, correlation phylogenetics, etc.). Moreover, the transfer efficiency over time of the stored OMVs was evaluated. The transfer rates remained unchanged for up to 10 days. Thereafter, the number of transformants gradually decreased for up to 30 days. Similar trends were shown in the study conducted by Chatterjee et al. on OMVs derived from *A. baumannii*, confirming the long-lasting stability without cryopreservatives [26]. Subsequently, it was investigated whether plasmid identity affected incorporation and transfer rate. The transfer of two different plasmids via *K. pneumoniae*-OMVs was examined, showing that the plasmid type induced changes in packaging and transformation rate. The high copy number plasmid (pGR) was loaded and transferred with greater efficiency compared to the low copy number plasmid (PRM). Our results were in line with a study conducted by Tran and Boedicker, in which the low copy number plasmid (pZS2501) had a low loading capacity (0.49 × 10^3^ copies per pg of OMVs), while the high-copy number plasmids (pLC291 and pUC19) showed a high loading potential (2.58 × 10^3^ and 482.7 × 10^3^ copies per pg of OMVs) [16]. Therefore, the plasmid cargo in the OMVs was strictly dependent on the copy number: the higher the PCN, the greater the plasmid amount in the OMVs and consequently the transformation efficiency. OMVs-mediated transfer exceeds the limits observed in other HGT-mechanisms [39]. Chatterjee et al. have already reported the ability of *A. baumannii*-OMVs to allow interspecies gene transfer [26]. For this reason, interspecies gene exchange was observed via *K. pneumoniae*-OMVs, using 4 different recipient species: *E. coli*, *S. enterica*, *P. aeruginosa* and *B. cepacia*. The generalized transfer to the different bacterial genera highlighted the HGT-OMVs efficiency, which verified independently of the phylogenetic correlation between the donor and recipient cell. Our experimental evidence showed that OMVs contributed to genetic exchange in microbial communities even among distantly related bacteria, without specific exchange mechanisms. Future studies will examine the possibility of OMVs to exchange DNA between different Gram-positive species.

In summary, the present study demonstrates, for the first time, the resistance gene to β-lactams spreads through OMVs secreted by *K. pneumoniae*. This innovative HGT mechanism allows for intra-species or inter-species diffusion, persistent over time and apparently not associated with specific limitations. Our future objectives will be studies aimed at blocking vesicular biogenesis, particularly of multidrug-resistant strains, to limit the spread of antibiotic resistance.

## Materials and Methods

### Bacterial Strains, plasmids and growing conditions

The strains used in this study were obtained from the American Type Culture Collection (ATCC) (Manassas, USA). *K. pneumoniae* ATCC 10031 was used for the OMVs purification. *K. pneumoniae* was transformed using the calcium chloride method with pGR (*K. pneumoniae*-pGR) (Addgene, Massachusetts, USA) and PRM-GFP (*K. pneumoniae*-PRM) (Addgene, Massachusetts, USA) respectively [40,41]. The first one was a high copy number plasmid (500 ~ 600 copies) containing genes for green fluorescent protein (GFP) and β-lactamase which conferred resistance to ampicillin. PRM was a plasmid containing the same genes and differed in copy number (10 ~ 12 copies). After transformation, *K. pneumoniae*-pGR and *K. pneumoniae*-PRM were cultured on Luria-Bertani agar (LB) (Sigma-Aldrich, St. Louis, USA) containing 100 μg mL^−1^ of ampicillin (Sigma-Aldrich, St. Louis, USA). *E. coli* ATCC 25922, *S. enterica* ATCC 14028, *P. aeruginosa* ATCC 13388 and *Burkholderia cepacia* ATCC 25416 were used as recipient strains for the HGT mediated by OMVs. All bacterial strains were cultured in LB (Sigma-Aldrich, St. Louis, USA) medium at 37 °C under orbital shaking at 180 rpm.

### OMVs Purification

OMVs were isolated from liquid cultures of *K. pneumoniae*-pGR and *K. pneumoniae*-PRM-GFP as previously described with modifications [42]. Ten milliliters of overnight (O/N) bacterial culture were inoculated in 600 mL of LB containing 100 μg mL^-1^ ampicillin. The bacterial inoculum was cultured at 37 °C under orbital shaking (180 rpm) for 8-12 hours, up to the OD_600_ nm value of 1. The cultures were centrifuged at 4000 × g at 4 °C for 20 min, to remove bacterial cells. Supernatants were decanted and filtered using vacuum Stericup^™^ 0.45 μm and 0.22 μm pore size polyethersulfone (PES) top filter (Millipore, Massachusetts, USA), to deflect remaining bacteria and cell debris. Vesicles were collected from cell-free supernatant culture by ultracentrifugation at 100000 × g (centrifuge Optima XPN-100 Beckman Coulter and rotor SW28) at 4 °C for 1.5 hours. Pellets were washed in sterile phosphate buffered saline 1X (PBS) by ultracentrifugation (100.000 × g at 4 °C for 1.5 hours). Vesicular pellets were suspended in 250 μL of PBS 1X and OMVs sterility was checked by inoculating 10 μL of vesicles on LB agar plates. OMV samples were treated with DNase (Applied Biological Materials – abm, British Columbia, Canada) according to the manufacturer’s protocol and stored at −20 °C until use.

### Transmission electron microscopy (TEM)

Purified OMVs were visualized by TEM, using negative staining. Five microliters of sample were adsorbed on carbon-coated copper/palladium grids for 30 min. A drop of sterile deionized water was used to wash the grids and a negative staining was realized by addition of 5 μL of 1% (w / v) uranyl acetate. TEM images were acquired using an EM 208 S transmission electron microscope (Philips, Amsterdam, Netherlands).

### OMVs size characterization by Dynamic Light Scattering (DLS)

Vesicles diameter size (Z-ave) and PDI analysis were performed using Zetasizer Nano-ZS (Malvern Instruments, Worcestershire, UK). For DLS, 40 μL of OMVs aliquot were mixed gently and transferred to sterile cuvettes. All measurements were conducted at 25 °C and three independent experiments for each purification were performed. DLS data were processed using Zetasizer software (V 7.11) provided by Malvern Panalytical (Malvern, UK).

### OMVs protein profile by tandem mass spectrometry (MS/MS)

For protein profile, OMVs were incubated with 1% Triton X-100 for 1 h at 4 °C. Lysed vesicles were centrifuged at 14.000 × g at 4 °C for 30 min and the supernatant was examined for protein amount by Bradford assay (HIMEDIA, Maharashtra, India). The protein extract was subjected to 10% sodium dodecyl sulfate polyacrylamide gel electrophoresis (SDS-PAGE). The gel was stained with Coomassie Brilliant blue G250 (Sigma-Aldrich St. Louis, USA) and different bands were cut to perform MS and MS / MS analysis, as previously described [14]. Briefly, protein bands were extracted from the gel and digested with trypsin. NanoUPLC-hr MS / MS analysis of the resulting peptide mixtures were performed on a Q-Exactive orbitrap mass spectrometer (Thermo Fisher Scientific, USA), coupled with a nanoUltimate300 UHPLC system (Thermo Fisher Scientific, USA). For protein identification, mass spectra were subjected to analysis by Mascot software (v2.5, Matrix Science, Boston, MA, USA), using the non-database redundant UniprotKB / Swiss-Prot (version 2020_03). The identified proteins were analyzed by subcellular localization, biological processes and molecular functions using Uniprot software (https://www.uniprot.org/).

### Intra-vesicular DNA analysis

Plasmid concentration in OMVs was determined by Real-time PCR (qPCR) using BrightGreen qPCR MasterMix Kits (abm, British Columbia, Canada), according to the manufacturer’s instructions. For DNA extraction, vesicles were lysed by boiling at 100 °C for 10 minutes. Two microliters of OMVs were added to 0.2 μM of primer, 1X mastermix in a final reaction volume of 20 μL. Primers used for qPCR were: β-lactamase Fw 5’- AACTTTATCCGCCTCCATCC-3’, β-lactamase Rev 3’- GCTATGTGGCGCGGTATTAT-5’. The amplification was performed in CFX96 Touch Real-Time PCR Detection System (Bio-Rad, California, USA), using the following amplification program: denaturation at 95 °C for 15 second, annealing at 60 °C for 20 second and extension at 72 °C for 15 second (40 cycles). The standard curves were constructed using purified plasmids from *K. pneumoniae*-pGR and *K. pneumoniae*-PRM respectively. Plasmid concentration in OMVs was converted in plasmid copy number (PCN), according to the formula:

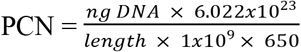

(http://cels.uri.edu/gsc/cndna.html). Subsequently, plasmid loading was estimated based on OMVs protein concentrations.

### OMVs mediated gene transfer

For gene transfer experiments through OMVs, the recipient strains *K. pneumoniae* ATCC, *E. coli* ATCC, *P. aeruginosa* ATCC, *B. cepacia* ATCC and *S. enterica* ATCC were inoculated in LB-broth up to OD_600_ nm value of 0.4. Cells were diluted in cold LB at final concentration of 10^7^ CFU / mL. Bacterial suspensions (60 μL) were incubated with 10 μg of OMVs statically for 4 hours at 37 °C and subsequently, for 4 hours under orbital shaking (180 rpm) at 37 °C. Fresh LB medium was added to each bacterial suspension and then incubated O/N under orbital shaking (180 rpm) at 37 °C. To further confirm that the plasmid transfer was mediated by OMVs, two separate experiments were performed with: (i) free plasmid and (ii) untreated cells. The following day, a 100 μL aliquot of bacteria was plated on LB-agar supplemented with 100 μg mL^-1^ ampicillin and incubated O/N at 37°C. The bacterial colonies (C_1-8_) were counted to define the transformation efficiency, according to the formula:

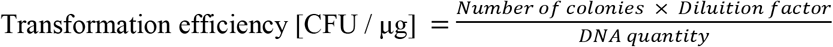

The same transformation experiments were performed using OMVs stored at −20 ° C for 10, 20 and 30 days and OMVs stored at +4°C for 3 and 7 days.

### Polymerase Chain Reaction (PCR) screening

After OMVs gene transfer, bacterial colonies grown on LB-agar supplemented with 100 μg mL^−1^ ampicillin were selected and subjected to molecular investigation for the presence of the plasmid by colony-PCR. Each bacterial colony was lysed by heat-shock and then centrifuged at 16.000 × g at 4°C for 10 min. The supernatant was transferred to a new Eppendorf and the DNA concentration was examined by NanoDrop 1000 spectrophotometer (Thermo Fisher Scientific, Massachusetts, USA). PCR were performed in a total volume of 50 μL containing 1 μM each primer, 1X Taq Master Mix (abm, British Columbia, Canada) and 100 ng of DNA. The primers used to amplify the 424 bp region of β-lactamase gene were Fw 5’- AACTTTATCCGCCTCCATCC-3’, Rev 3’- GCTATGTGGCGCGGTATTAT-5’. The amplification was conducted in Thermal cycler UNO96 (VWR International, Pennsylvania, USA) according to the following program: initial denaturation at 94°C for 3 min, 35 cycles of amplification in which each cycle was denatured at 94 °C for 30 second, annealed at 57.3 °C for 30 second and extended at 72 °C 1 min; the final extension at 72 °C for 5 min. As a housekeeping gene control, 16S rRNA gene was amplified, using the primers: Fw 5’- GGTAGAGTTTGATCCTGGCTCAG-3’, Rev 3’- ATTACCGCGGCTGCTGG-5’. The used program was: initial denaturation at 94°C for 1 min, 30 cycles of amplification in which each cycle was denatured at 94 °C for 1 min, annealed at 58 °C for 1 min and extended at 72 °C 1.5 min; the final extension at 72 °C for 10 min. To visualize the amplification product, 1% agarose gel electrophoresis was performed.

### Antibiotic susceptibility test

The disk diffusion assay was performed according to the National Committee on Clinical Laboratory Standards (NCCLS) [43]. Fresh colonies, before and after OMVs treatment, were inoculated in physiological solutions to 0.5 McFarland turbidity. With cotton swab dipped in the bacterial inoculum, the solution was homogeneously plated into Mueller-Hinton (MH) agar plates. Disks of ampicillin (10 μg) (Thermo Fisher Scientific, Massachusetts, USA), piperacillin (30 μg) (Thermo Fisher Scientific, Massachusetts, USA) and ciprofloxacin (5 μg) (Thermo Fisher Scientific, Massachusetts, USA) were placed on the plates and were incubated at 37 °C O/N. The antibiotic susceptibility was examined by measuring the zone of inhibition diameter, according to the European Committee on Antimicrobial Susceptibility Testing (EUCAST) guidelines.

## Author Contributions

Conceptualization: Federica Dell’Annunziata

Data curation: Veronica Folliero, Anna De Filippis

Formal analysis: Giuliana Donadio, Nunzianna Doti

Funding acquisition: Giovanni Boccia

Investigation: Carmela Dell’Aversana

Project administration: Massimiliano Galdiero

Supervision: Gianluigi Franci

Validation: Viviana Izzo

Writing – original draft: Marilena Galdiero, Lucia Altucci

Writing – review & editing: Fabrizio Dal Piaz

## References

1. Bedhomme S, Amorós-Moya D, Valero LM, Bonifaci N, Pujana M-À, Bravo IG. Evolutionary Changes after Translational Challenges Imposed by Horizontal Gene Transfer. Genome Biol Evol. 2019;11: 814–831. doi:10.1093/gbe/evz031

2. Bello-López JM, Cabrero-Martínez OA, Ibáñez-Cervantes G, Hernández-Cortez C, Pelcastre-Rodríguez LI, Gonzalez-Avila LU, et al. Horizontal Gene Transfer and Its Association with Antibiotic Resistance in the Genus Aeromonas spp. Microorganisms. 2019;7. doi:10.3390/microorganisms7090363

3. Emamalipour M, Seidi K, Zununi Vahed S, Jahanban-Esfahlan A, Jaymand M, Majdi H, et al. Horizontal Gene Transfer: From Evolutionary Flexibility to Disease Progression. Front Cell Dev Biol. 2020;8: 229. doi:10.3389/fcell.2020.00229

4. Davies J, Davies D. Origins and evolution of antibiotic resistance. Microbiol Mol Biol Rev. 2010;74: 417–433. doi:10.1128/MMBR.00016-10

5. Hall RJ, Whelan FJ, McInerney JO, Ou Y, Domingo-Sananes MR. Horizontal Gene Transfer as a Source of Conflict and Cooperation in Prokaryotes. Front Microbiol. 2020;11: 1569. doi:10.3389/fmicb.2020.01569

6. Ely B. Recombination and gene loss occur simultaneously during bacterial horizontal gene transfer. PLoS One. 2020;15: e0227987. doi:10.1371/journal.pone.0227987

7. Redondo-Salvo S, Fernández-López R, Ruiz R, Vielva L, de Toro M, Rocha EPC, et al. Pathways for horizontal gene transfer in bacteria revealed by a global map of their plasmids. Nat Commun. 2020;11: 3602. doi:10.1038/s41467-020-17278-2

8. Riva F, Riva V, Eckert EM, Colinas N, Di Cesare A, Borin S, et al. An Environmental Escherichia coli Strain Is Naturally Competent to Acquire Exogenous DNA. Front Microbiol. 2020;11: 574301. doi:10.3389/fmicb.2020.574301

9. Salvadori G, Junges R, Morrison DA, Petersen FC. Competence in Streptococcus pneumoniae and Close Commensal Relatives: Mechanisms and Implications. Front Cell Infect Microbiol. 2019;9: 94. doi:10.3389/fcimb.2019.00094

10. Headd B, Bradford SA. The Conjugation Window in an Escherichia coli K-12 Strain with an IncFII Plasmid. Appl Environ Microbiol. 2020;86. doi:10.1128/AEM.00948-20

11. Grohmann E, Muth G, Espinosa M. Conjugative plasmid transfer in gram-positive bacteria. Microbiol Mol Biol Rev. 2003;67: 277–301, table of contents. doi:10.1128/MMBR.67.2.277-301.2003

12. Fillol-Salom A, Alsaadi A, Sousa JAM de, Zhong L, Foster KR, Rocha EPC, et al. Bacteriophages benefit from generalized transduction. PLoS Pathog. 2019;15: e1007888. doi:10.1371/journal.ppat.1007888

13. Gómez-Gómez C, Blanco-Picazo P, Brown-Jaque M, Quirós P, Rodríguez-Rubio L, Cerdà-Cuellar M, et al. Infectious phage particles packaging antibiotic resistance genes found in meat products and chicken feces. Sci Rep. 2019;9: 13281. doi:10.1038/s41598-019-49898-0

14. Dell’Annunziata F, Ilisso CP, Dell’Aversana C, Greco G, Coppola A, Martora F, et al. Outer Membrane Vesicles Derived from Klebsiella pneumoniae Influence the miRNA Expression Profile in Human Bronchial Epithelial BEAS-2B Cells. Microorganisms. 2020;8. doi:10.3390/microorganisms8121985

15. Domingues S, Nielsen KM. Membrane vesicles and horizontal gene transfer in prokaryotes. Curr Opin Microbiol. 2017;38: 16–21. doi:10.1016/j.mib.2017.03.012

16. Tran F, Boedicker JQ. Genetic cargo and bacterial species set the rate of vesicle-mediated horizontal gene transfer. Sci Rep. 2017;7: 8813. doi:10.1038/s41598-017-07447-7

17. Fulsundar S, Harms K, Flaten GE, Johnsen PJ, Chopade BA, Nielsen KM. Gene transfer potential of outer membrane vesicles of Acinetobacter baylyi and effects of stress on vesiculation. Appl Environ Microbiol. 2014;80: 3469–3483. doi:10.1128/AEM.04248-13

18. Wang S, Gao J, Wang Z. Outer membrane vesicles for vaccination and targeted drug delivery. Wiley Interdiscip Rev Nanomed Nanobiotechnol. 2019;11: e1523. doi: 10.1002/wnan.1523

19. Yoon H. Bacterial Outer Membrane Vesicles as a Delivery System for Virulence Regulation. J Microbiol Biotechnol. 2016;26: 1343–1347. doi:10.4014/jmb.1604.04080

20. Schwechheimer C, Kuehn MJ. Outer-membrane vesicles from Gram-negative bacteria: biogenesis and functions. Nat Rev Microbiol. 2015;13: 605–619. doi:10.1038/nrmicro3525

21. Turnbull L, Toyofuku M, Hynen AL, Kurosawa M, Pessi G, Petty NK, et al. Explosive cell lysis as a mechanism for the biogenesis of bacterial membrane vesicles and biofilms. Nat Commun. 2016;7: 11220. doi:10.1038/ncomms11220

22. Cecil JD, Sirisaengtaksin N, O’Brien-Simpson NM, Krachler AM. Outer Membrane Vesicle-Host Cell Interactions. Microbiol Spectr. 2019;7. doi:10.1128/microbiolspec.PSIB-0001-2018

23. Caruana JC, Walper SA. Bacterial Membrane Vesicles as Mediators of Microbe - Microbe and Microbe - Host Community Interactions. Front Microbiol. 2020;11:432. doi:10.3389/fmicb.2020.00432

24. Verhoeven AJ, Estrela JM, Meijer AJ. Alpha-adrenergic stimulation of glutamine metabolism in isolated rat hepatocytes. Biochem J. 1985;230: 457–463. doi:10.1042/bj2300457

25. Kolling GL, Matthews KR. Export of virulence genes and Shiga toxin by membrane vesicles of Escherichia coli O157:H7. Appl Environ Microbiol. 1999;65: 1843–1848. doi:10.1128/AEM.65.5.1843-1848.1999

26. Chatterjee S, Mondal A, Mitra S, Basu S. Acinetobacter baumannii transfers the blaNDM-1 gene via outer membrane vesicles. J Antimicrob Chemother. 2017;72: 2201–2207. doi:10.1093/jac/dkx131

27. Blesa A, Berenguer J. Contribution of vesicle-protected extracellular DNA to horizontal gene transfer in Thermus spp. Int Microbiol. 2015;18: 177–187. doi:10.2436/20.1501.01.248

28. Ho M-H, Chen C-H, Goodwin JS, Wang B-Y, Xie H. Functional Advantages of Porphyromonas gingivalis Vesicles. PLoS One. 2015;10: e0123448. doi:10.1371/journal.pone.0123448

29. Yaron S, Kolling GL, Simon L, Matthews KR. Vesicle-mediated transfer of virulence genes from Escherichia coli O157:H7 to other enteric bacteria. Appl Environ Microbiol. 2000;66: 4414–4420. doi:10.1128/AEM.66.10.4414-4420.2000

30. Rumbo C, Fernández-Moreira E, Merino M, Poza M, Mendez JA, Soares NC, et al. Horizontal transfer of the OXA-24 carbapenemase gene via outer membrane vesicles: a new mechanism of dissemination of carbapenem resistance genes in Acinetobacter baumannii. Antimicrob Agents Chemother. 2011;55: 3084–3090. doi:10.1128/AAC.00929-10

31. Navon-Venezia S, Kondratyeva K, Carattoli A. Klebsiella pneumoniae: a major worldwide source and shuttle for antibiotic resistance. FEMS Microbiol Rev. 2017;41: 252–275. doi:10.1093/femsre/fux013

32. Lopes E, Saavedra MJ, Costa E, de Lencastre H, Poirel L, Aires-de-Sousa M. Epidemiology of carbapenemase-producing Klebsiella pneumoniae in northern Portugal: Predominance of KPC-2 and OXA-48. J Glob Antimicrob Resist. 2020;22: 349–353. doi:10.1016/j.jgar.2020.04.007

33. Martora F, Pinto F, Folliero V, Cammarota M, Dell’Annunziata F, Squillaci G, et al. Isolation, characterization and analysis of pro-inflammatory potential of Klebsiella pneumoniae outer membrane vesicles. Microbial Pathogenesis. 2019;136: 103719. doi:10.1016/j.micpath.2019.103719

34. Lerner A, Matthias T, Aminov R. Potential Effects of Horizontal Gene Exchange in the Human Gut. Front Immunol. 2017;8: 1630. doi:10.3389/fimmu.2017.01630

35. Lerminiaux NA, Cameron ADS. Horizontal transfer of antibiotic resistance genes in clinical environments. Can J Microbiol. 2019;65: 34–44. doi:10.1139/cjm-2018-0275

36. Da Silva GJ, Domingues S. Insights on the Horizontal Gene Transfer of Carbapenemase Determinants in the Opportunistic Pathogen Acinetobacter baumannii. Microorganisms. 2016;4. doi:10.3390/microorganisms4030029

37. Ha JY, Choi S-Y, Lee JH, Hong S-H, Lee H-J. Delivery of Periodontopathogenic Extracellular Vesicles to Brain Monocytes and Microglial IL-6 Promotion by RNA Cargo. Front Mol Biosci. 2020;7: 596366. doi: 10.3389/fmolb.2020.596366

38. Pérez-Cruz C, Carrión O, Delgado L, Martinez G, López-Iglesias C, Mercade E. New type of outer membrane vesicle produced by the Gram-negative bacterium Shewanella vesiculosa M7T: implications for DNA content. Appl Environ Microbiol. 2013;79:1874–1881. doi:10.1128/AEM.03657-12

39. Aminov RI. Horizontal gene exchange in environmental microbiota. Front Microbiol. 2011;2: 158. doi:10.3389/fmicb.2011.00158

40. Chan W-T, Verma CS, Lane DP, Gan SK-E. A comparison and optimization of methods and factors affecting the transformation of Escherichia coli. Biosci Rep. 2013;33: e00086. doi:10.1042/BSR20130098

41. Higuchi-Takeuchi M, Morisaki K, Numata K. Method for the facile transformation of marine purple photosynthetic bacteria using chemically competent cells. MicrobiologyOpen. 2020;9. doi:10.1002/mbo3.953

42. De Lise F, Mensitieri F, Rusciano G, Dal Piaz F, Forte G, Di Lorenzo F, et al. Novosphingobium sp. PP1Y as a novel source of outer membrane vesicles. J Microbiol. 2019;57: 498–508. doi:10.1007/s12275-019-8483-2

43. Petrillo F, Pignataro D, Lavano MA, Santella B, Folliero V, Zannella C, et al. Current Evidence on the Ocular Surface Microbiota and Related Diseases. Microorganisms. 2020;8. doi: 10.3390/microorganisms8071033

